# Overcoming sparseness of biomedical networks to identify drug repositioning candidates

**DOI:** 10.1101/2020.06.07.138966

**Authors:** Aleksandar Poleksic

## Abstract

Modeling complex biological systems is necessary to understand biochemical interactions behind pharmacological effects of drugs. Successful *in silico* drug repurposing requires a thorough exploration of diverse biochemical concepts and their relationships, including drug’s adverse reactions, drug targets, disease symptoms, as well as disease associated genes and their pathways, to name a few. We present a computational method for inferring drug-disease associations from complex but incomplete and biased biological networks. Our method employs the compressed sensing technique to overcome the sparseness of biomedical data and, in turn, to enrich the set of verified relationships between different biomedical entities. We present a strategy for identifying network paths supportive of drug efficacy as well as a computational procedure capable of combining different network patterns to better distinguish treatments from non-treatments. The data and programs are freely available at http://bioinfo.cs.uni.edu/AEONET.html.

## 1 Introduction

Recent decades have seen a sharp increase in spending on drug development in US and the rest of the world. The number of drugs approved per 1 billion dollars invested in R&D has steadily declined since 1950^1^, with the estimates of the average cost to bring a new drug to the market approaching 1.7 billion ^2^. Reversing the trend of the stagnant and diminishing drug approval rates requires adoption of new approaches to drug discovery. To address the toxicity and lack of efficacy of candidate drugs, researchers are increasingly looking for therapeutics that act on multiple targets pertaining to single or multiple disease pathways. Such a broader approach to the development of pharmaceutical agents, coined as polypharmacology^3,4^, often necessitates the use of multiple chemicals acting on different targets for the treatment of complex diseases. Polypharmacology is the main principle behind drug repurposing, which aims to identify new uses for the existing drugs. Drug repurposing accelerates the discovery process by bypassing most of the pre-clinical work and trials required for drug approval.

Computational methods provide crucial support for drug repurposing. Modern algorithms increasingly rely on the network representation of a biological system to discover drug-disease, drug-gene, disease-gene, and other relationships between biomedical concepts. A simple biological system can be viewed as a homogenous network in which the nodes represent biological entities and edges represent the relationships (associations or interactions) between those entities. An example is the protein interaction network, in which the nodes represent proteins and edges signify PPIs.

While homogenous network can be used to model relationships between entities from any pair of domains, the accumulating scientific evidence suggests that better understanding of a biological system as a whole is necessary to overcome the limited performance of methods that only take account of one or two types of biological data. Complex biological systems can be viewed as heterogeneous networks in which the nodes are grouped into different layers (domains), such as the compound layer, protein layer, disease layer, etc. The nodes within each layer are interconnected and connected to the nodes from other layers^5^.

Heterogeneous networks provide new avenues for research into *in silico* drug repurposing^6,7,8,9^. Zeng *et al*. used a multi-modal autoencoder on a heterogeneous network to learn drug features and, in turn, prioritize drug repurposing candidates^10^. Chen *et al*. developed an efficient algorithm for detecting missing cross-layer links in a heterogeneous network by leveraging the observed cross-layer dependency with the within-layer topology^11^. Guney *et al*. developed a drug-disease proximity measure to quantify the network-based associations between drugs and disease proteins^12^. Cheng *et al*. proposed a systems pharmacology-based procedure that integrates molecular pharmacoepidemiologic and network-based approaches to quantify interplay between drug targets and disease proteins^13^.

Perhaps the most comprehensive biological network to date is HetioNet^14^. This heterogeneous network, compiled from public resources^15–54^, consists of 47,031 nodes grouped into 11 domains and 2,250,197 relationships (edges) classified into 24 categories (Fig. 1).

**Fig 1.**
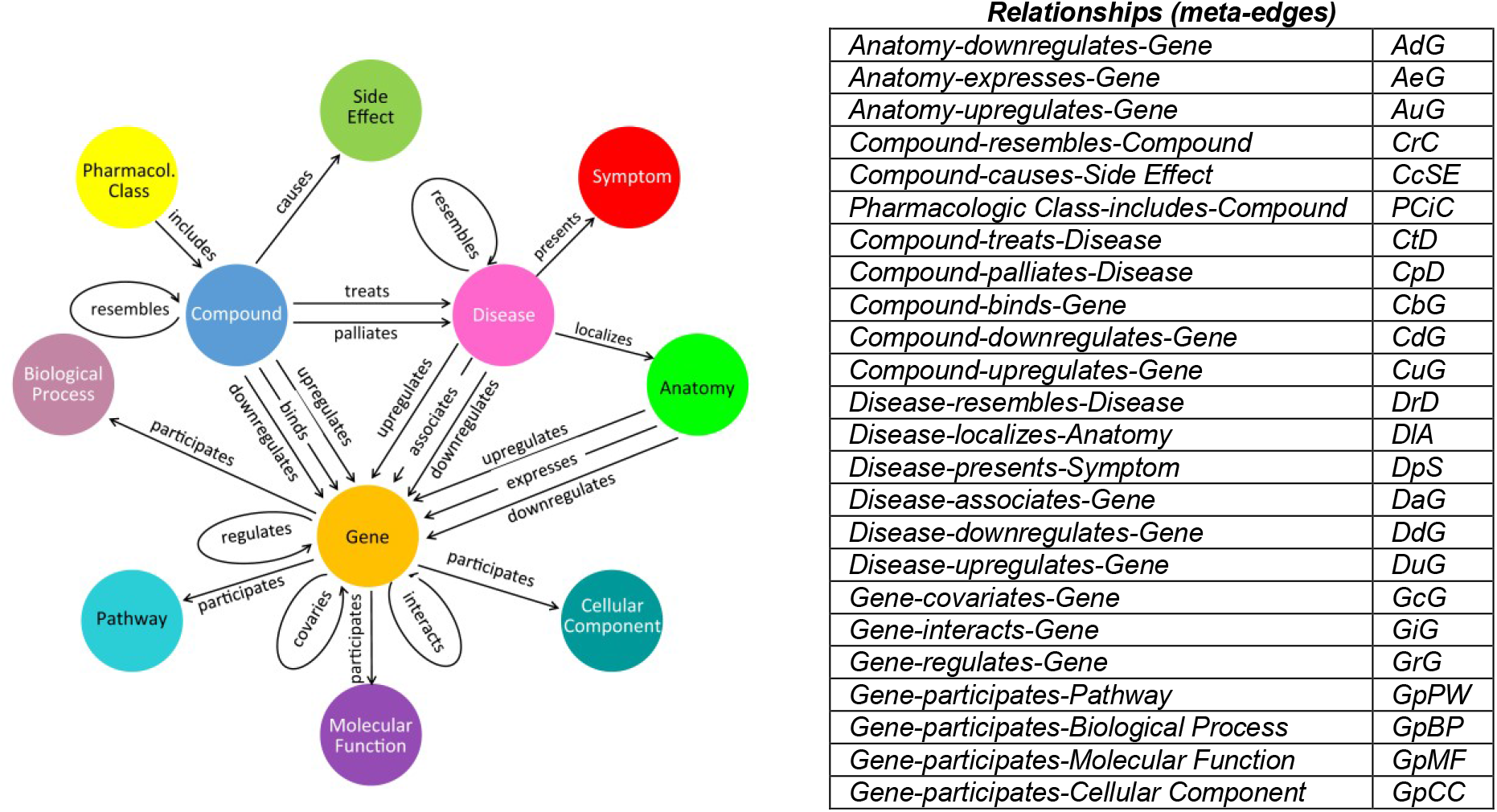
HetioNet

In the project Rephetio, Himmelstein *et al*. demonstrated the advantage of utilizing HetioNet patterns to reconstruct known and predict new drug-disease associations^14^. Rephetio is capable of identifying multiple network paths that explain therapeutic efficacy of different drugs on different diseases. The method uses logistic regression to fine-tune the weights of different network paths that connect compounds to diseases, including transitive paths that traverse multiple relations, such as, for example, *Compound-binds–Gene–participates–Pathway–participates–Gene– associates–Disease*. (The meta-path in this example represents any path from a compound node *c* to a disease node *d* with the property that *c* binds a gene *g_1_* which participates in the same pathway as gene *g_2_*, which is associated to the disease *d*.) The aggregate probability over selected paths is used to measure the strength of each drug-disease association.

While the novel network-based approach used by Himmelstein *et al*. is useful in gaining insight into the drug mechanism of action, the accuracy of Rephetio is highly dependent on the available network data. Here we present a computational method AeoNet, which is capable of identifying network patterns that explain drug efficacy in context of sparse, biased and incomplete networks. Our algorithm can be viewed as a two-step process.

First, we use the compressed sensing technique ^55,56,57^ to enrich the set of network edges by inferring all missing inter-domain and cross-domain relationships. In the second step, we integrate the likelihoods of all possible paths connecting each drug with each disease. We use three external benchmarks to demonstrate that our approach consistently increases the accuracy of drug-disease association prediction.

The rest of this paper is organized as follows. In section 2.1 we provide a brief overview of compressed sensing. Section 2.2 describes a simple procedure for constructing different network paths that connect drugs to diseases. Section 2.3 presents a strategy for selecting a set of the most informative paths, while section 2.4 presents a simple method for aggregating drug-disease association probabilities along those paths. The Results section summarizes the performance of our method on the external validation sets.

## 2 Methods

### 2.1 Inferring missing HetioNet relationships

We use a variant of the matrix completion technique, referred to as “compressed sensing”, to enrich the set of edges between any two HetioNet domains connected by a meta-edge. Compressed sensing is similar in spirit to dual regularized one-class collaborative filtering (OCCF) algorithm^58^. However, in contrast to OCCF, our method is based on logistic matrix factorization^59^ and thus has a more explicit statistical foundation.

#### The compressed sensing algorithm

In what follows, we illustrate how this procedure works in an example setting of drug-target interaction prediction. The same algorithm is applied to predict relationships between entities that belong to other directly connected biological domains.

There are two scientific premises behind our method. First, we assume that the matrix of drug-target association (that we wish to recover) is of small rank *k* ≪ min{*n*, *m*}, where *n* and *m* denote the total number of drugs and genes, respectively. Our algorithm (Fig. 2) can be viewed as the low-rank completion of the input (noisy and incomplete) drug-target association matrix *R* = (*r*_*i,j*_), defined as

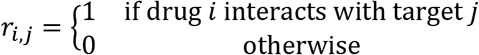

**Fig. 2.**
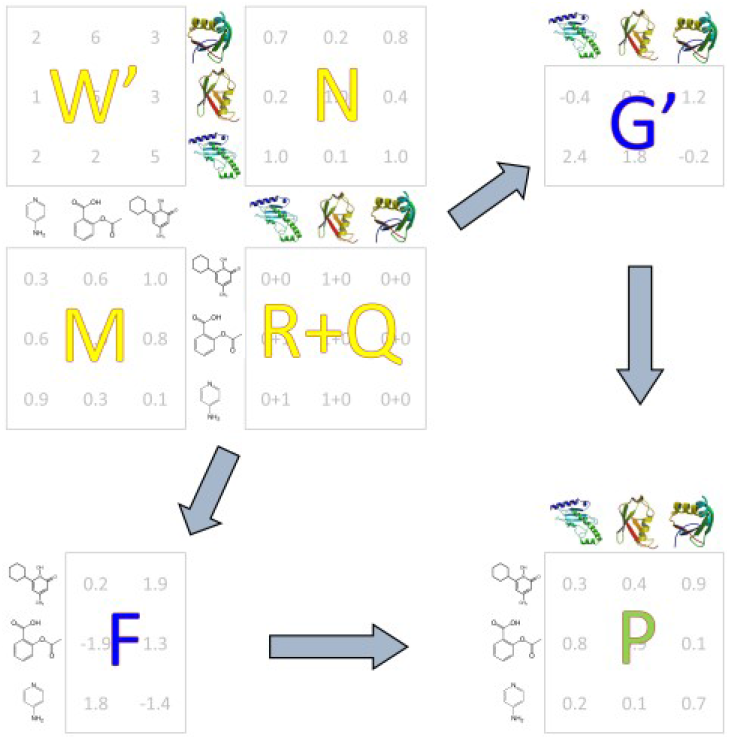
Compressed sensing algorithm. R: input matrix of drug-target associations; Q: impute matrix; W: weight matrix; M: drug similarity matrix; N: target similarity matrix; F: latent drug preferences; G: latent target preferences; P: output matrix of drug-target interaction probabilities.

Our second assumption is that similar drugs interact with similar targets. Given the relationship matrix *R*, the pairwise drug similarity matrix *M*, and the pairwise target similarity matrix *N*, the algorithm computes the matrices *F* and *G* of drugs’ and targets’ “latent” preferences (Fig. 2). Specifically, the *i*^*th*^ row of *F* represents the latent vector representation of the drug *i* while the *i*^*th*^ row of *G* represents the latent vector representation of target *i*. The matrices *F* and *G* are found by minimizing the loss function

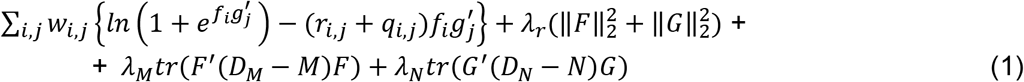

In (1) above, the prime symbol denotes matrix transpose, ‖ ‖_2_ stands for the Frobenius norm, and *tr* denotes the matrix trace. We use *f_i_* and *g_j_* to denote the *i*^*th*^ row of *F* and *j*^*th*^ row of *G*, respectively. The matrix *D*_*M*_ is the diagonal “degree matrix” of *M* in which the element *d_i,i_* represents the sum of the elements in the *i*^*th*^ row of *M*. The (optional) impute values *q_i,j_* can be specified by expert users who wish to supplement the set of relationships *R*, while the entries *w_i,j_* of the weight matrix *W* reflect the user’s confidence in the elements of the interaction matrix *R* + *Q*. The lambdas (*λ_r_*, *λ_M_*, and *λ_N_*) are optimizable parameters. The matrix of probabilities of drug-target interactions is computed as *P* = exp(*FG*′) /(1 + exp(*FG*′)). The latent preferences of drugs with no known interacting targets and targets with no known interacting drugs are derived with a variant of the weighted profile method. For technical details, we refer the reader to our published work^55,56,57^.

### 2.2 Exploring different paths connecting drugs to diseases

We compute drug-disease association probabilities separately along each path between the drug layer and the disease layer.

The variety of paths between two layers gives rise to a set of diverse similarity matrices *M* and *N* that can be specified as input to our compressed sensing algorithm. To derive side information specific to a particular network path *p*, we split *p* into three sub-paths, *p_1_*, *p_2_*, and *p_3_*. The sub-path *p_1_* is a cycle that starts and ends at the first layer and uniquely defines a similarity matrix *M*. The sub-path *p_2_* consists of a single edge that connects the first layer to the second layer and corresponds to the relationship itself (matrix *R*). Finally, the sub-path *p_3_*, determining the side information *N*, is a cycle that starts and ends at the second layer.

For instance, the path *p* = *CrCtDrD* (*Compound-resembles-Compound-treats-Disease-resembles-Disease*) can be represented as *p* = *p_1_p_2_p_3_*, where *p_1_* = *CrC*, *p_2_* = *CtD*, and *p_3_* = *DrD*. The drug-disease association probabilities along this path are derived by running the compressed sensing algorithm on the input drug similarity matrix *M* = *CrC* (*Compound-resembles-Compound*), the association matrix *R* = *CtD* (*Compound-treats-Disease*), and the disease similarity matrix *N* = *DrD* (*Disease-resembles-Disease*). The same idea is extended to derive *M*, *R*, and *N* along longer paths. For instance, as illustrated in Fig. 3, the drug-disease probabilities along the path *CcSEcCtDpSpD* (*Compound-causes-Side Effect-causes-Compound-treats-Disease-causes-Symptom-causes-Disease*) are computed by running the compressed sensing algorithm on the input matrices *M* = *CcSEcC*, *R* = *CtD*, and *N* = *DpSpD*, where *CcSEcC* (abbreviated as *CseC*) represents the matrix of pairwise similarities of drugs’ side-effect profiles and *DpSpD* (or short *DpsD*) represents the matrix of pairwise similarities of disease’ symptom profiles (Table 1, Fig. 3). More specifically, given compounds *i* and *j*, we set the element of the matrix *CseC* at the position (*i*, *j*) to the Jaccard similarity of the rows *i* and *j* of the *CcSE* matrix (*Compound-causes-Side Effect*). The disease similarity matrix *N* = *DpsD* is specified in a similar manner, by computing Jaccard similarities of diseases’ symptom profiles stored in the relationship matrix *DpS* (*Disease-presents-Symptom*).

**Fig. 3.**
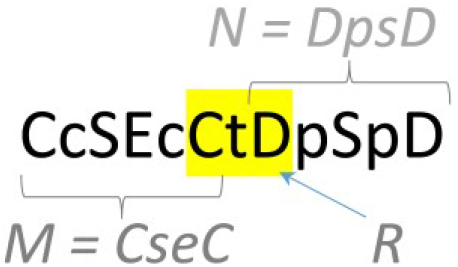
Assigning path-dependent matrices *R*, *M* and *N*.

**Table 1.** Different drug, gene, and disease similarity matrices.

| Compound Similarity Matrix | Derived from HetioNet Relationship |
| --- | --- |
| CrC | Compound-resembles-Compound |
| CseC | Compound-causes-Side Effect (CcSE) |
| CpcC | Pharmacologic Class-includes-Compound (PCiC) |
| CsC | Average of CseC and CpcC |
| Disease Similarity Matrix | Derived from HetioNet Relationship |
| DrD | Disease-resembles-Disease |
| DpsD | Disease-presents-Symptom (DpS) |
| DlaD | Anatomy (DIA) |
| DsD | Average of DpsD and DlaD |
| Gene Similarity Matrix | Derived from HetioNet Relationship |
| GbpG | Biological Process (GpBP) |
| GmfG | Molecular Function (GpMF) |
| GccG | Cellular Component (GpCC) |
| GpwG | Pathway (GpPW) |
| GsG | Average of GbpG, GmfG, GccG, GpwG, GiG, GrG, GcG |

In theory, *p_1_* or *p_3_* (or both) can be of length zero, indicating the lack of side information. In those cases, the input similarity matrices are simply set to the identity matrices. For instance, if *p* = *CrCtD*, then *M* = *CrC*, *R* = CtD, and *N* = *I* (no drug side information).

Finally, we define the probabilities of drug efficacies along transitive paths as the normalized products of the probability matrices for individual path segments. For instance, the matrix of drug-disease associations along the path *CseCbGiGaDpsD* (*Compound-causes-Side Effect-causes-Compound-binds-Gene-interacts-Gene-associates-Disease-presents-Symptom-presents-Disease*) represents the normalized product of the probability matrices computed along the paths *CseCbGiG and GiGaDpsD*.

While multiple network patterns provide different clues into therapeutic effects of different drugs on different diseases, accounting for all possible paths is computationally expensive (see section 2.4). One way to lower the cost of our algorithm is to merge selected paths into paths that traverse new edges, those corresponding to the overall drug, disease and gene similarity relationships (denoted by *CsC*, *DsD*, and *GsG*, respectively). We define the mean drug similarity score as the average of the similarity scores based on side-effect and pharmacological profiles, or, at the matrix level, *CsC* = (*CseC* + *CpcC*) / 2. In a similar way, we compute the average similarity of diseases as *DsD* = (*DpsD* + *DlaD*) / *2* and the average similarity of genes as *GsG* = (*GbpG* + *GpwG* + *GmfG* + *GccG* + *GiG* + *GcG* + *GrG*) / 7. The notion of average similarity helps reduce the total number of paths explored by AeoNet. For instance, using the average compound similarity measure, the paths *CseCtDlaD* and *CpcCtDlaD* are merged into a single path *CsCtDlaD*. Table 2 shows the collection of all paths through the network explored by AeoNet.

**Table 2.**
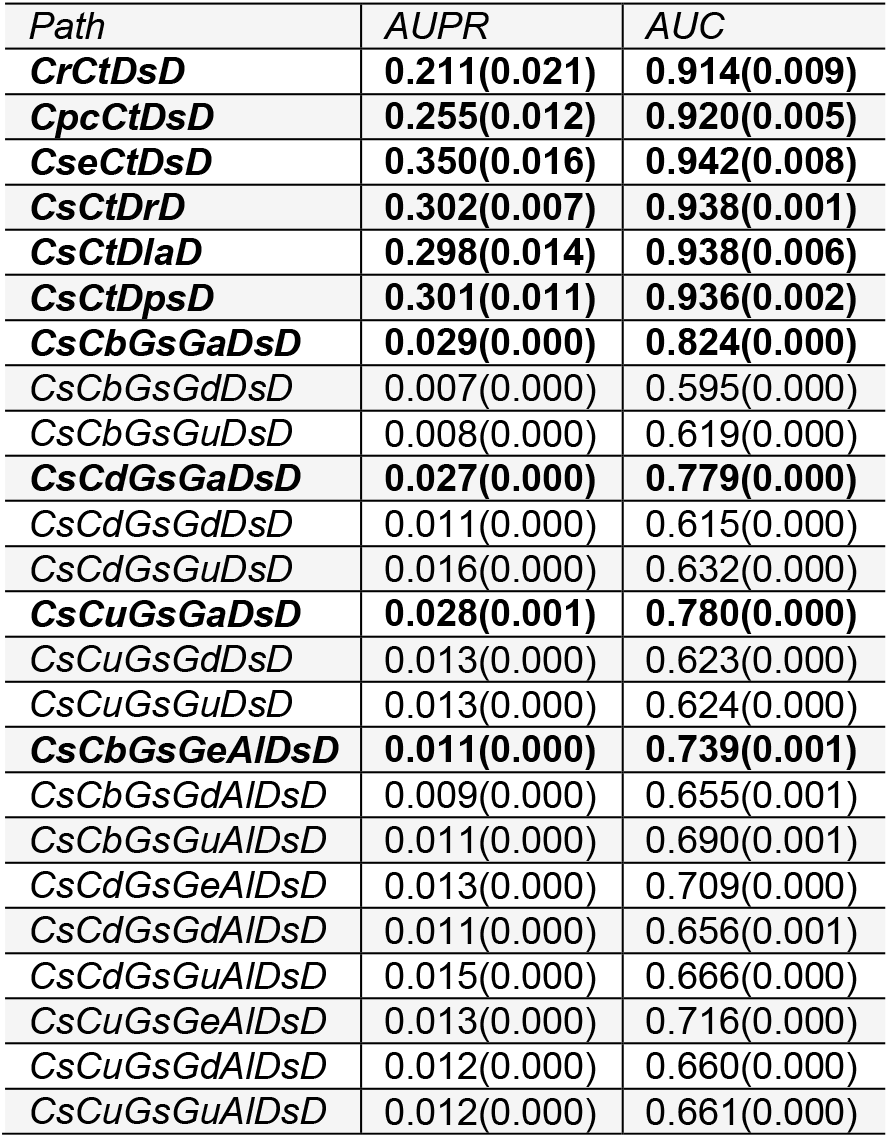
Classification accuracy of different network patterns (three rounds of 3-fold CV). The paths with highest AUC scores are shown in bold.

### 2.3 Measuring contribution of paths supporting drug efficacy

Different network paths have different potentials in distinguishing disease treatments from non-treatments. In general, we have less confidence in drug-disease association scores computed along longer paths or paths that traverse meta-edges derived from sparse databases of cross-layer relationships. The treatment potential of different network patterns is computed using cross validation on the set of HetioNet drug-disease relationships (*Compound-treats-Disease*). Table 2 shows cross-validated AUC, AUPR and PREC@10 scores corresponding to different network patterns.

### 2.4 Aggregating drug-disease probabilities along the most informative paths

AeoNet integrates probabilities along the network paths with highest cross-validated AUC (Table 2, Fig. 4). We note that nine out of ten highest scoring AUC paths also represent the highest scoring paths with respect to the AUPR measure, indicating a good agreement between the two classification metrics on the set of 755 indications (*Compound-treats-Disease*).

**Fig. 4.**
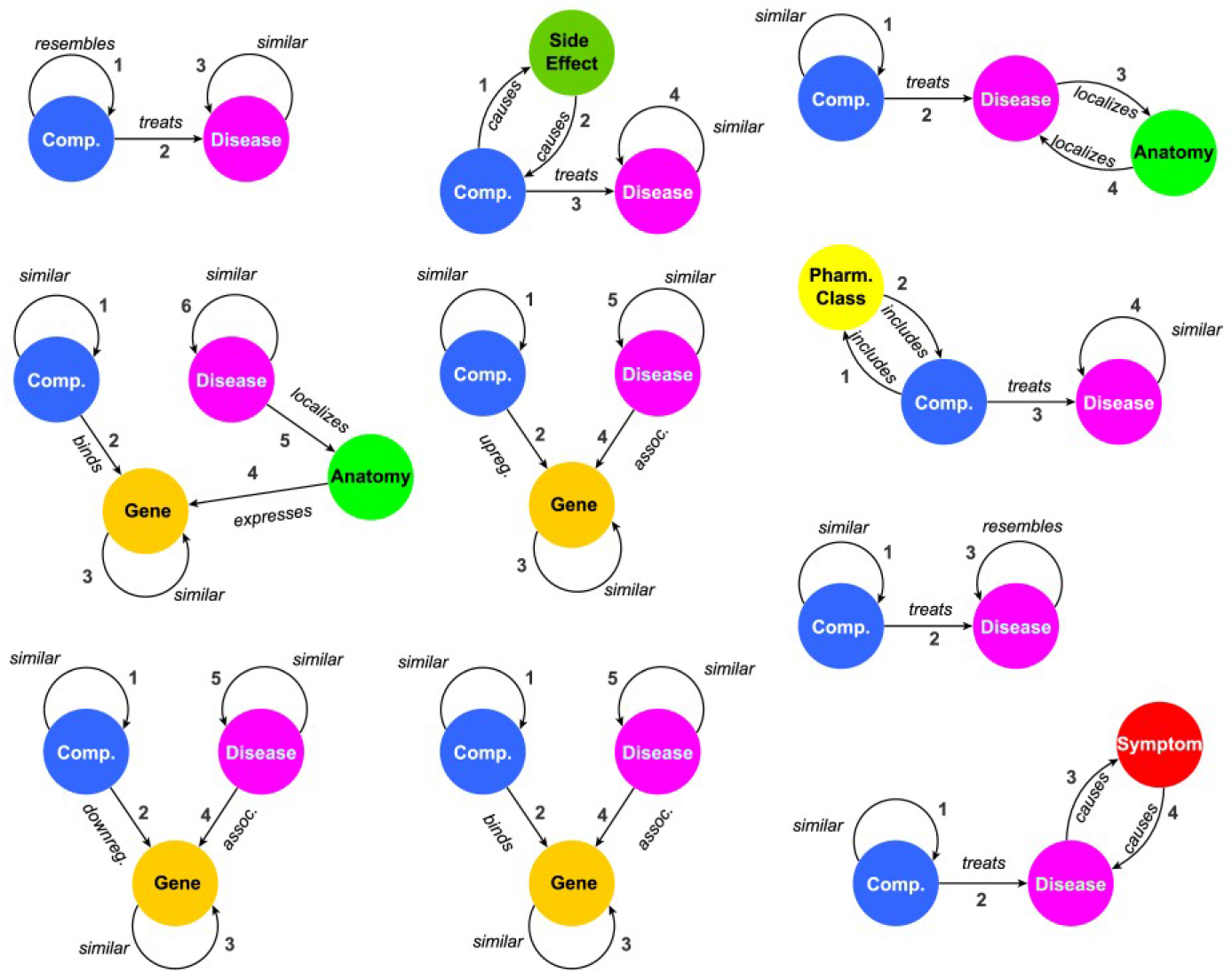
Most supportive network paths (in no particular order) as measured by cross-validated AUC score.

An optimal weighted sum of drug-disease probabilities along ten highest scoring paths is found by maximizing cross-validated AUC score achieved on the set of *Compound-treats-Disease* associations. The path coefficients are selected from *C* = {3^0^, 3^1^, 3^2^, 3^3^} using grid search. More specifically, for each among 4^10^ permutations with repetitions of 10 coefficients selected from *C*, we compute cross-validated AUC of the weighted sum of path specific probabilities and then pick the set of 10 coefficients that yields the highest AUC.

While a larger selection of network paths combined with a finer grid search has potential to improve the method’s accuracy further, we had to scale both numbers down in order to balance the computational cost with the available computing resources. For predicting palliative (as opposed to disease-modifying) potentials of drugs, we exclude paths that traverse *Compound-treats-Disease* edges.

## 3 Results

We assess the potential of different network paths to distinguish treatments from non-treatments and the overall accuracy of AeoNe using three external validation sets compiled by Himmelstein *et al*., namely DrugCentral, Clinical Trials and Symptomatic. The DrugCentral test set consists of 208 novel treatments and 207,572 non-treatments extracted from the DrugCentral repository^40^. The Clinical Trial set consists of 5,594 novel indications compiled from ClinicalTrials.gov and 202,186 non-treatments. We note that, while much larger, the Clinical Trial benchmark is less reliable than DrugCentral, due to the low approval rates of drugs in clinical trials. The Symptomatic set contains 390 HetioNet relationships (positive associations) of the type *Compound-palliates-Disease*^14^ and 208,023 negative associations. We emphasize that no positive drug-disease association from any test sets appears in our training set of 755 HetioNet disease-modifying indications (*Compound-treats-Disease* relationships).

Table 3 list the classification scores along the individual network paths and the scores obtained using the integrative AeoNet approach. As seen in this table, the value added by different network paths varies across the three test sets. For instance, the path with the highest AUPR score in the DrugCentral benchmark is *CpcCtDsD* (AUPR = 0.095) but this path does not perform nearly as well in other benchmarks (it ranks #6 in DrugCentral AUC benchmark and #6 and #8 in Clinical Trial AUPR and AUC benchmarks, respectively). On the other hand, the path *CsCbGsGaDsD* has the highest AUC score (AUC = 0.747) and the second highest AUPR score (AUPR = 0.097) in Clinical Trial benchmark but it ranks in the bottom half of all paths in the DrugCentral benchmarks with respect to both AUC and AUPR. By finding and exploiting synergies between different network paths, AeoNet consistently improves classification scores of individual network paths (Table 3).

**Table 3.**
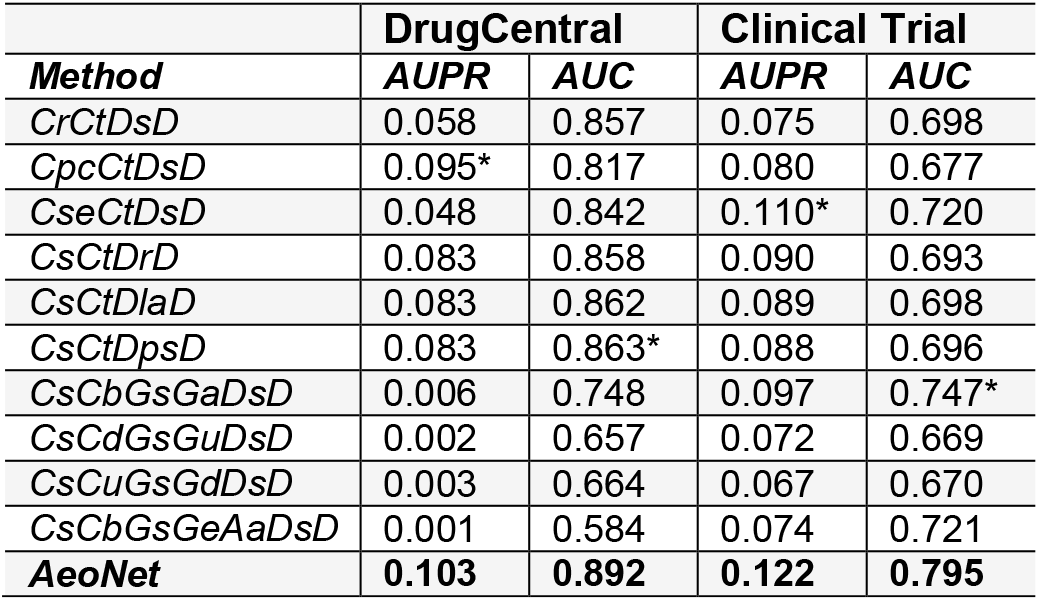
Classification accuracy along individual network paths and the accuracy of the integrative approach. Highest scoring individual paths are denoted by *. Best method overall is shown in bold.

We also present a side-by-side comparison of AeoNet with Rephetio and two other recently published methods for predicting drug-associated indications, BNNR^60^ and OMC^61^. The last two algorithms have been shown to yield higher accuracy when compared to other state-of-the-art approaches. These methods require drug similarity information in terms of Tanimoto coefficients as well as the disease similarity information. We computed Tanimoto coefficients by calculating Jaccard similarity between the drug fingerprints derived from the Chemistry Development Kit, using PaDEL-Descriptor software^62^. For disease side information, both BNNR and OMC were given *Disease-resembles-Disease* scores. In addition, the OMC method tested here is based on tri-layer network (OMC3), and it requires drug-gene and disease-gene association matrices as input. We derived this data from HetioNet’s *Drug-binds-Gene* and *Disease-associates-Gene* relationships, respectively. All three external methods were ran using the default parameters supplied by their authors.

With the exception of Clinical Trial PREC@10 test (OMC 40%, AeoNet 40%), the classification scores achieved by AeoNet exceed the scores achieved by the other methods in every benchmark and according to each classification measure (Table 3, Fig. 5). The performance of the remaining three methods fluctuate across the test sets and across different benchmarking measures used. For instance, Rephetio has the second highest AUC scores in every benchmark but all other methods compare favorably to Rephetio in PREC@10 tests. The AUC scores achieved by OMC trail those achieved by AeoNet and Rephetio but this method shows a consistently good performance in AUPR and PREC@10 tests.

**Fig. 5.**
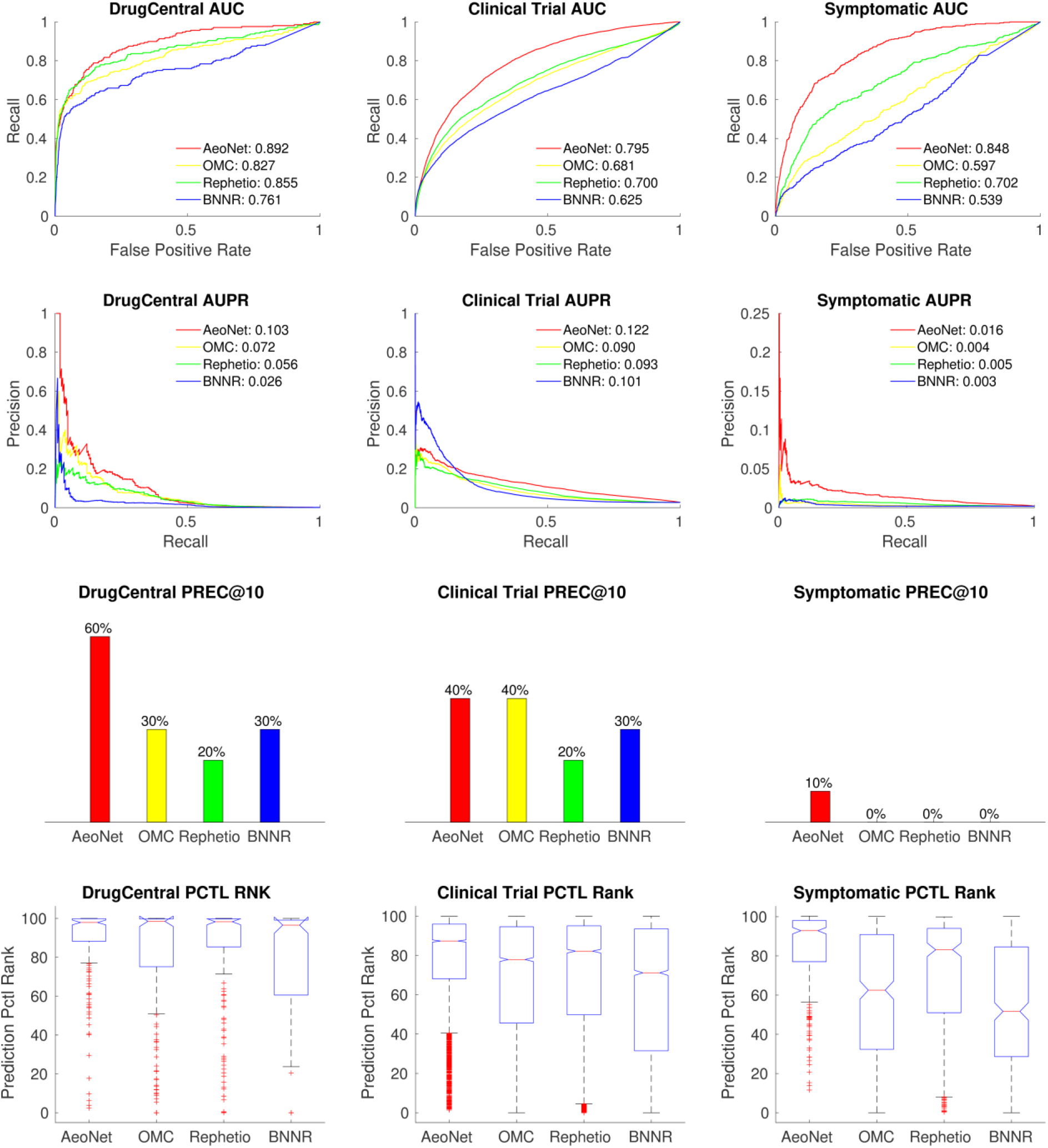
Comparing AeoNet with state-of-art using different classification metrics: AUPR, AUC, PREC@10, and Prediction Percentile Rank.

The superior performance of our method in recovering Symptomatic relationships is suggestive of different action mechanisms of disease-modifying therapies compared to palliative ones. We re-emphasize that AeoNet computes probabilities of palliative therapies along the transitive paths from drugs to diseases (those that pass through the gene nodes) ignoring all paths that include Compound-treats-Disease edge.

The seemingly low AUPR scores of the four methods in the three benchmark need proper interpretation. To place the AUPR scores in context, we compare them to the scores obtained by a purely random classifier. The AUPR score achieved by a random classifier is equal to the fraction of condition positives in the test set (∑ *cond. pos* / (∑ *cond. pos* + ∑ *cond. neg*)). For instance, the AUPR score achieved by a random classifier on DrugCentral data set is ~0.001 (208 condition positives and 207,572 condition negatives). For comparison, on the same data set the AeoNet method achieves the AUPR score of 0.103, which represents two orders of magnitude (100 fold) enrichment over the random classifier.

We observe that, at any p-value cutoff, the fraction of DrugCentral indications recovered by AeoNet is much larger compared to the percentage of recovered Clinical Trial indications (Fig 6). This is expected and desirable, as a majority of Clinical Trial drugs fail to get FDA approval. In fact, the recovery rate of Clinical Trial indications exceeding the FDA drug approval rate (currently around 9.6%^63^) should be indicative of poor method’s specificity.

**Fig. 6.**
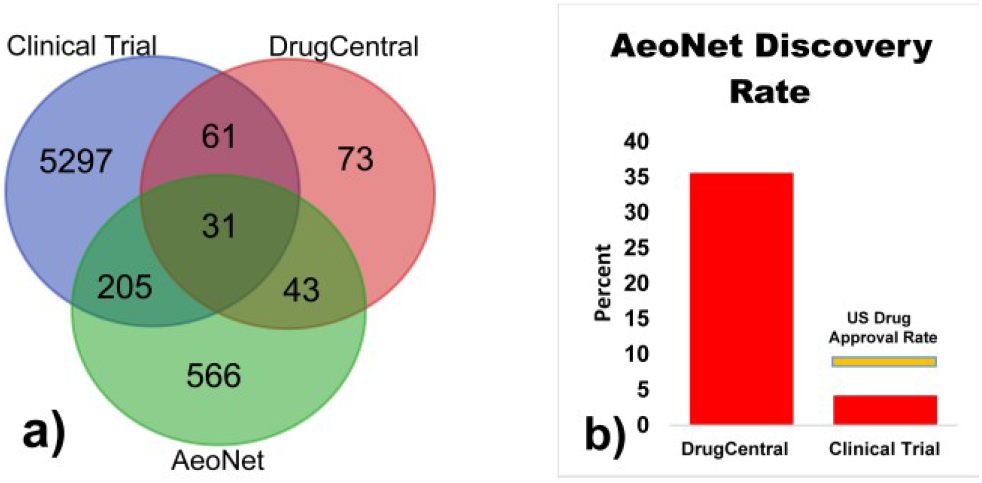
(a) The overlap of 845 AeoNet predictions with p-values ≤1.0E-3 with those stored in DrugCentral and Clinical Trial (training set indications not included) (b) Percentage of DrugCentral and Clinical Trial indications found by AeoNet using the same p-vaue cutoff. Current FDA drug approval rate is shown in orange.

## Illustrative example – Bone Cancer

Dactinomycin is an intravenous therapy indicated for Ewing’s sarcoma, a type of cancer that grows in bone or surrounding tissues, mostly in children and young adults. Dactinomycin represents a good repurposing test for AeoNet, as its association with bone-cancer is listed in DrugCentral but not in HetioNet.

The top six AeoNet predictions for the treatment of bone cancer are Epirubicin, Cisplatin, Methotrexate, Carboplatin, Doxorubicin, and Dactinomycin (in ranked order). We note that the first five hits are relatively easy cases for AeoNet, because the associations of those drugs with bone-cancer are already recorded in HetioNet. AeoNet places Dactinomycin (p-value=2.2E-8) on top of all drugs that are not listed as one of 755 HetioNet disease-modifying therapies for bone-cancer, despite the fact that the drug lacks structural analogs among drugs with similar indications in HetioNet. In fact, the closest structural neighbor of Dactinomycin in HetioNet, as measured by Tanimoto coefficient, is Moxifloxacin, an antibiotic used to treat bacterial infections. Moreover, a closer inspection of *Compound-resembles-Compound* relationships reveals that no drug “resembles” Dactinomycin in HetioNet, further illustrating AeoNet’s ability to make accurate inferences based on aggregate probabilities of different transitive network paths.

## Illustrative example – Gout

Gout is a general term for different conditions caused by the deposits of monosodium urate crystals, typically in the joints of the foot or ankle. There are six treatments of gout in DrugCentral that are not present in HetioNet. We wanted to find out whether AeoNet can confidently predict any of those six drugs as potential repurposing candidates.

After excluding gout associations recorded in HetioNet (which are consistently found in the highest ranks by our method) the top eight AeoNet predictions for gout treatments include four drugs listed in DrugCentral but not HetioNet, namely Triamcinolone, Dexamethasone, Betamethasone and Azathioprine. While the first three hits have p-values below 1.0E-13 and despite these drugs not being present in our training set, we still consider them relatively easy predictions, as each one of those drugs has a close chemical analog in HetioNet. However, Azathioprine (p-value = 2.0E-4) is a difficult prediction case since a closer inspection of *Compound-resembles-Compound* relationships reveals that Azathioprine does not resemble any drug with known HetioNet indications. Moreover, Azathioprine is not structurally similar to any other gout treatment in HetioNet (Tanimoto coef. < 0.27). Fig. 7 shows three strong Azathioprine-goat supporting paths identified by AeoNet, including a path that span two novel edges that do not appear in HetioNet, namely *Azathioprine upregulates-SPP1* (predicted p-value: 4.3E-12) and *gout-associates-SPP1* (p-value: 8.9E-16)^64^.

**Fig. 7.**
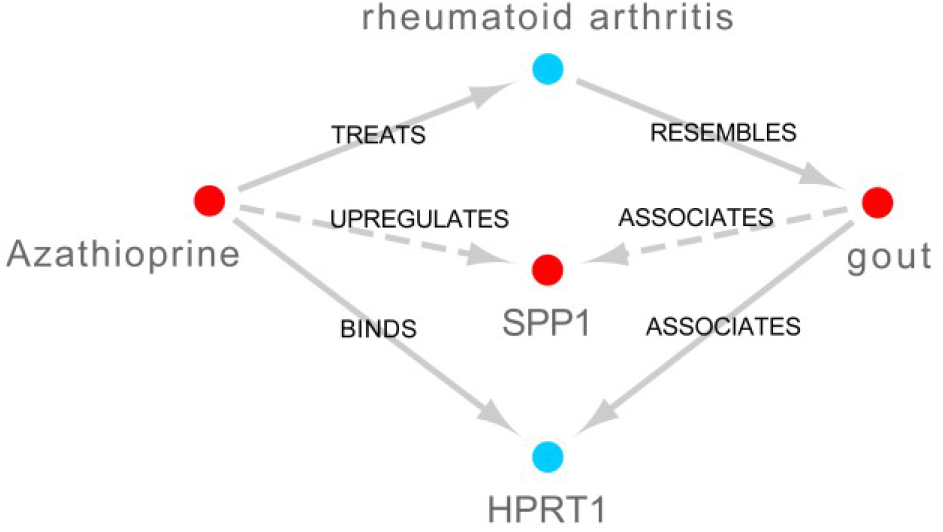
Most significant paths of length 2 connecting Azathioprine and gout. Solid edges represent existing HetioNet relationships. Dashed edges (forming a new network path) are predicted by compressed sensing.

## Discussion and Conclusion

Heterogeneous networks are emerging as a powerful way to model complex biological systems. Network implementations allow the inference of relationships among distantly related biological entities and improve the prediction of associations between closely related entities. We present a computational technique for finding potential drug repurposing candidates from a comprehensive network of interactions between tens of thousands of biomedical entities, including drugs, side-effects, diseases, symptoms, and genes. Our methodology for distinguishing disease treatments from non-treatments integrates the likelihoods of different network patterns supporting drug efficacy. In contrast to previous studies that seek to explore only experimentally verified relationships, the method we propose operates on a much larger, probabilistic network that consists of tens of millions of predicted relationships between the network nodes. With minor modifications, our algorithm can be used in other inference tasks, such as drug-target interaction or disease-gene association prediction.

The results of our study hint at some straightforward network enhancement that can further increase the prediction accuracy. For instance, expanding the network to include compound structural similarity, gene homology and/or gene-side effect associations^65^ would enrich the set of paths supporting drug efficacy and, in turn, provide further insights into drug mechanisms of actions. These modifications, along with the fast growing biomedical databases and advances in machine learning algorithms will fuel research into computational drug discovery and repurposing in the years to come.

## Acknowledgements

We are thankful to Prof. Lei Xie from CUNY Graduate Center for inspiring discussions on the topic. We also thank Daniel Himmelstein for helping us compile the three external validation sets. This work is supported in part by the University of Northern Iowa Professional Development award, the Capacity Building grant and the Google Cloud grant.

## Notes

http://bioinfo.cs.uni.edu/AEONET.html

### Competing Interest Statement

The authors have declared no competing interest.

